# Dynamic and Depth Dependent Nanomechanical Properties of Dorsal Ruffles in Live Cells and Biopolymeric Hydrogels

**DOI:** 10.1101/189852

**Authors:** Varun Vyas, Melani Solomon, Gerard G. M. D’Souza, Bryan D. Huey

## Abstract

The nanomechanical properties of various biological and cellular surfaces are increasingly investigated with Scanning Probe Microscopy. Surface stiffness measurements are currently being used to define metastatic properties of various cancerous cell lines and other related biological tissues. Here we present a unique methodology to understand depth dependent nanomechanical variations in stiffness in biopolymers and live cells. In this study we have used A2780 & NIH3T3 cell lines and 0.5% & 1% Agarose to investigate depth dependent stiffness and porosity on nanomechanical properties in different biological systems. This analytical methodology can circumvent the issue associated with the contribution of substrates on cell stiffness. Here we demonstrate that by calculating ‘continuous-step-wise-modulus’ on force vs. distance curves one can observe minute variation as function of depth. Due to the presence of different kinds of cytoskeletal filament, dissipation of contact force might vary from one portion of a cell to another. On NIH3T3 cell lines, stiffness profile of Circular Dorsal Ruffles could be observed in form of large parabolic feature with changes in stiffness at different depth. In biopolymers like agarose, depending upon the extent of polymerization in there can be increase or decrease in stiffness due variations in pore size and extent to which crosslinking is taking place at different depths. 0.5% agarose showed gradual decrease in stiffness whereas with 1% agarose there was slight increase in stiffness as one indents deeper into its surface.

## Introduction

Atomic Force Microscopy (AFM) is an important tool for nanomechanical investigations of cells and biomaterials^1-4^, in which a specimen is indented with spatial resolution to the level of nanoscale in order to measure and or map the local modulus or related properties. For instance, differences in stiffness have been identified for healthy versus cancer cells, dissimilarities in the brush layer of normal versus cancerous cells, nanoscale heterogeneity in bones and other nanoscale properties have been studied with AFM probes^5-7^. Typically, this is achieved by measuring forces experienced by an AFM probe upon indentation into a specimen (‘Force-distance curves’). Measured at one location, or more usefully in arrays to map local properties, straightforward elastic mechanics are generally applied such as the Hertzian or Sneddon models. Often, such results are additionally compared to idealized controls such as hydrogels of agarose or collagen due to their perceived mechanical similarity to the cytoplasm of a cell ^8-10^.

While such analyses are sufficient for general as well as local observations of cellular mechanical properties, coupled AFM and optical microscopy in fact reveal that during and/or following nanomechanical indentation there can be substantial deformation of the local and overall cellular volume^11^, shape ^12^, and/ or sub-structure ^11, 13^. Since the cytoplasm is generally 1.2-1.4 times as viscous as water^14^, Force-distance curves may also vary due to pororelastic effects. This has been reported with HeLa cervical cancer cells and Madin-Darby canine kidney (MDCK) cells^15, 16^, with significant influences from the rate of water efflux for the cytoplasmic networks. The presence of the underlying substrate cannot be overlooked either, generally causing a smooth increase in stiffness as the indenting probe approaches the substrate unless numerically accommodated for example by the Bottom Effect Cone Correction (BECC) method^17^. More actively, the structure and hence mechanical properties of living cells can dynamically adjust during the minutes to hours of typical AFM measurements, either due to ongoing, ‘natural’ remodeling of internal structures with a living cell, or in direct response to chemo/mechanical perturbation by the AFM probe ^18^.

Perhaps most important for nanomechanical investigations of cells, and mesoscopic materials in general, is that their structure is generally not at all homogeneous, instead exhibiting sometimes extensive structural hierarchies. For example, aside from obvious cellular components such as a nucleus or nucleoid, stresses applied to the apical surface of cells can be transmitted across the cell body to focal adhesion points in the basal plane^19, 20^. In particular, the network of actin stress fibers in cells can heterogeneously distribute locally applied stresses during AFM-based indentation experiments to distant, inhomogeneously distributed focal adhesion centers ^21^. This should result in local variations in mechanical properties for cells, in the lateral and normal directions. Equivalently, the ensemble mechanical properties of inanimate mesoscopic or hierarchical materials systems, such as many cell scaffold and tissues engineering systems, could cause strong variations in local mechanical properties depending on the distribution, spatially and in terms of stiffness, of interconnected features. Accordingly, this work employs depth dependent indentation profiles (‘DDIP’), in which successive segments of Force-distance curves are analyzed throughout each indentation, providing a ‘continuous-modulus’ capable of identifying mechanical variations in 3-dimensions. This uniquely allows the measurement, visualization, and assessment of local mechanical properties, including the associated error parameters, as a function of depth and position, specifically applied to living mammalian cells but generally applicable to any mesoscopic materials systems. Here investigations are limited to depth dependence nanomechanical properties in a live cell rather than the loading rate which know to influence measured stiffness ^22^. To understand how loading rate influences depth dependent modulus is topic of separate investigation.

### Background

The procedure and benefits of depth dependent indentation profiling are demonstrated with *in vitro* AFM-indentations of living cells cultured onto glass coverslips from two specific cell lines, human ovarian carcinomas (A2780) and mouse fibroblasts (NIH3T3). Identical measurements on drop coated gels from 0.5% and 1% Agarose solution are also considered for comparison to this broadly employed standard, as sketched in Figure 1A. Notwithstanding the aforementioned mechanical complexity of living cells, the modulus is analyzed obeying standard elastic mechanical models for straightforward comparison to the wide body of AFM-based indentation literature. For optimal spatial resolution of the underlying nano‐ to micro-scale features, commercial square pyramidal AFM tips are utilized, indented in square arrays of 128 by 128 points (essentially pixels) over areas up to 40 μm on a side.

**Figure 1.**
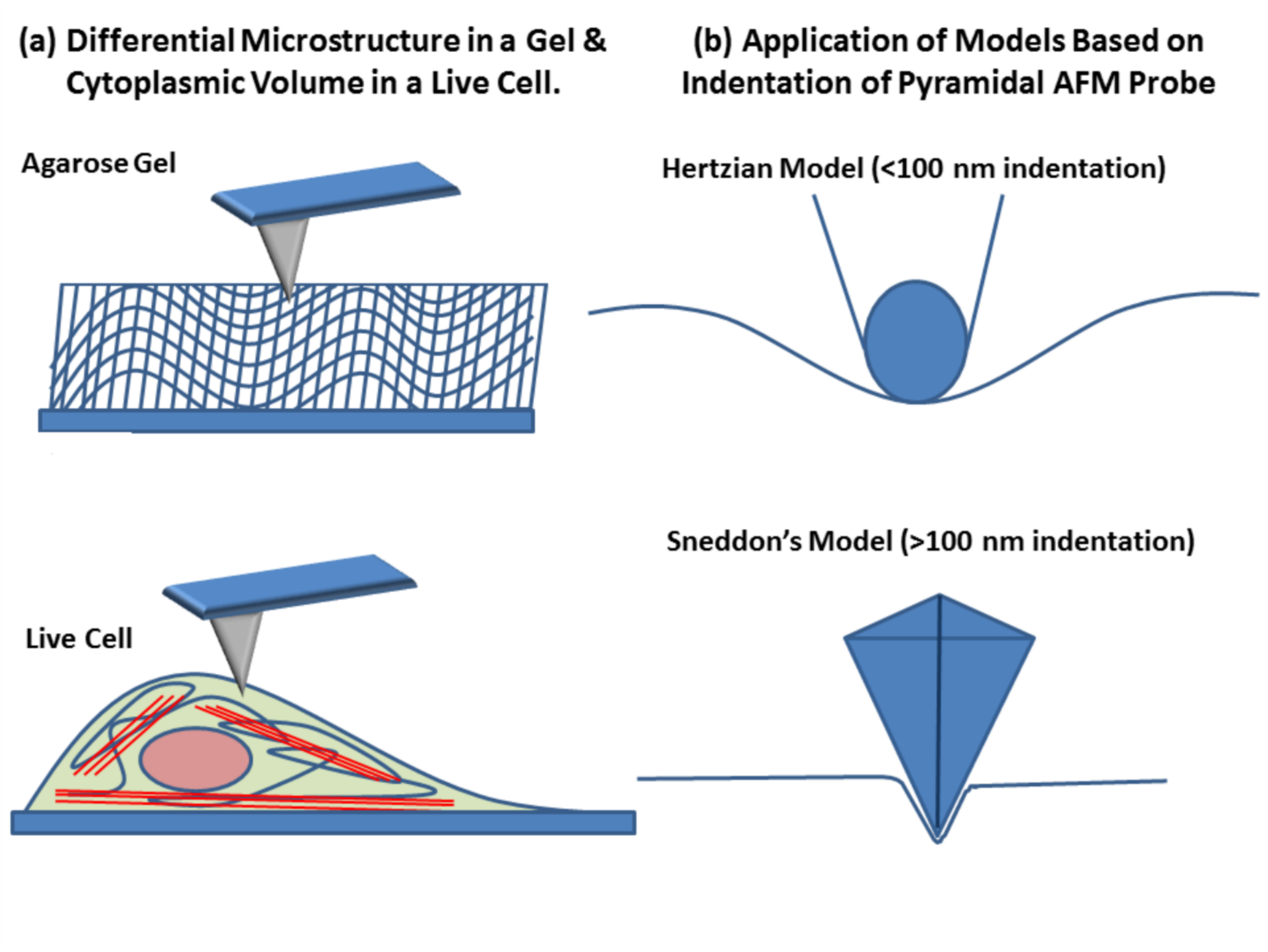
(a) Sketches of the microstructural variability in a gel (top row) or live cell (base) that may produce depth dependent variations in stiffness. (b) Appropriate mechanical models based on the indentation depth of the AFM probe.

Upon initial contact during each measurement, for indentation depths less than the diameter of the assumed sphere at the apex of the AFM probe (80 nm assumed here), the essentially hemispherical contact area is best described by Hertzian mechanics ^23^. As the probe indents deeper, however, contact between the deforming cell and the square-pyramidal geometry along the probe length will dominate, in which case Sneddon’s equation is more appropriate, Figure 1B. A general indentation equation is therefore presented (Equation 1) for interpreting the measured Force (F) versus indentation (δ) curves in either case. This includes a model dependent constant term (C_model_), the indentation to a model-dependent power (δ^n^), and the local reduced modulus (E*). These terms are specifically defined for Hertz (Equation 2) and Sneddon (Equation 3) mechanics to analyze initial contact (surface) and sub-surface stiffness, respectively^23, 24^. Standard definitions for the reduced radius (R*, Equation 4) and reduced modulus (E*, Equation 5) are also shown. For simplicity, the sample is assumed to be flat at the point of initial contact (infinite radius and hence the 2^nd^ term in Equation 4 drops out), while a poisson ratio (ν) of 0.3 is employed throughout for the sample. Note that the modulus (E) of the tip is so large (>100 GPa) compared to the sample (<1 MPa) that the corresponding term in Equation (5) is negligible; therefore, it may be ignored along with the poisson ratio of the tip in the final analysis as indicated.

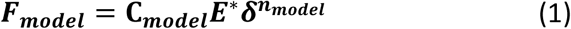

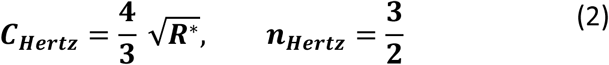

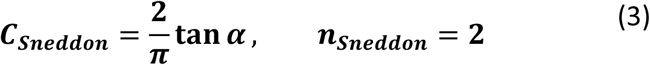

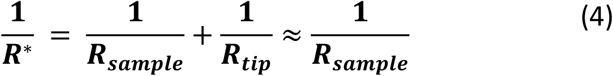

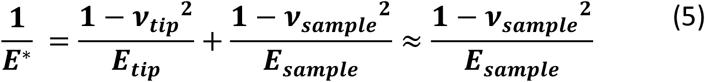

Ultimately, the local reduced modulus can finally be evaluated for any given Force-indentation curve, based on the slope of the n^th^ root of the Force versus the indentation, all taken to the n^th^ power, and divided by the appropriate constant (Equation 6). Beyond providing a generalized method to calculate the modulus, this approach also gives a straightforward assessment of the standard deviation of the modulus, seldom considered in the AFM literature. This is determined by substituting the standard deviation of the calculated slope, which is commonly employed in simple linear regression (Equation 7), in place of the actual slope of Equation 6 (in braces). The effective local sample modulus, and corresponding standard deviation of this measured local modulus, are finally determined for every pixel, and for every depth, using Equation 5. Far beyond implementing best scientific practices by incorporating such error analyses (however rarely it is reported), the error maps this provides are also found to be valuable in interpreting indentation results as uniquely shown in this work.

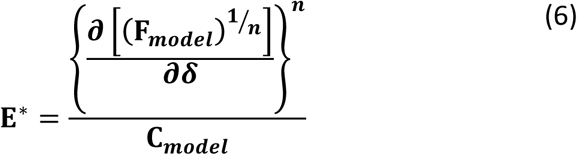

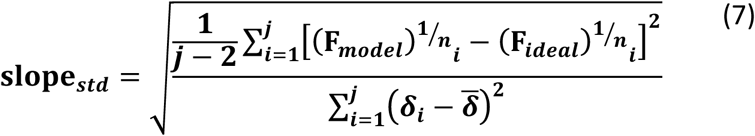

It is noted that similar approaches to the DDIP concept have been alluded in previous AFM studies of cellular mechanics^25, 26^, but extensive mapping of depth dependent nanomechanical properties hasn’t been a preferred choice for presenting and analyzing stiffness in cells and polymers. For example, a ‘point-wise-modulus’ was calculated for arrays up to 16x16 indentations by fitting to the Hertz model as deep as 400 nm, from which differences in the general behavior from standard Hertzian mechanics were interpreted as identifying regions with distinct composite-like properties ^27^. Depth dependent force profiles (instead of modulus) have also been widely considered, for instance recently to identify the density of polymer brushes at the surface of healthy versus cancer cells^6^. Here, DDIP is employed to quantify, and identify the depth of, any increase or decrease in the elastic stiffness as the probe indents deeper into a specimen, with error calculations evidencing non-elastic responses. This includes sub-surface structures, a dynamic response of the specimen (e.g. cytoskeletal assembly/disassembly), the underlying substrate, etc. Specifically, regions of varying effective stiffness are clearly mapped at different depths in living A2780 carcinoma cells. Circular dorsal ruffles (CDR)^28^, actin based membrane protrusions that have a critical role in receptor internalization and cell migration, are also believed to be identified in NIH3T3 mouse fibroblast cells. AFM-based measurements have not previously been reported of these dynamic cellular sub-structures, which exhibit residence times on the order of tens of minutes. Finally, equivalent measurements of Agar films, serving as inanimate, nearly homogeneous standards, provide guidance particularly in interpreting modulus error maps.

## Material and Methods

### Cell Culturing

Mouse fibroblast cells (NIH 3T3) and human ovarian carcinoma cells (A2780) were obtained from ATCC (Manassas, VA). The NIH3T3 culture was maintained in DMEM (Corning cellgro®, Manassas, VA) supplemented with 10% fetal bovine serum and 1% penicillin-streptomycin. A2780 cells were maintained in RPMI-1640 (Lonza, Alpharetta, GA) supplemented with 20% fetal bovine serum and 1% penicillin-streptomycin. Both cell lines were cultured in a 5% CO_2_, 37^°^C atmosphere. All AFM based measurements of these cells were performed in the same media.

### Agarose Film

10 ml of 0.5% or 1% agarose solutions were prepared in milli-q water and heated in a microwave for 3 mins per manufacturer instructions. This agarose solution was drop coated onto a glass coverslip. After cooling, force maps were acquired in milli-q water.

### Force Mapping Nanomechanical Measurements

AFM based force mapping measurements were conducted with an Asylum Research MFP-3D AFM with integrated inverted optics (Nikon TE-2000), using a liquid cell for complete immersion during the experiments. For cultured cell lines, arrays of 128 x 128 force indentations were acquired over a 40 μm x 40 μm area. For control measurements on drop coated agarose films, statistical relevance is more important than spatial resolution and hence multiple (3) 16 x 16 force maps over an area of 20 μm were collected for averaging and error calculations. All force maps were collected with indentation velocities of 32 μm/sec, amounting to individual indentations at a rate of ~2 Hz. Spring constants of the Olympus TR800PB probes used throughout were calibrated following the widely applied Sader method ^29^. All analysis, including maps of the Hertzian and Sneddon derived stiffness as a function of depth, as well as novel maps of the error in these calculations, were performed using custom analytical and visualization routines written with Matlab software.

## Result & Discussion

### Depth Dependent Indentation Profiles of A2780 human ovarian carcinomas

DDIP was performed on two distinct cell lines, NIH3T3 (Mouse Fibroblast) and A2780 (Human Ovarian Carcinomas), both of which have each been extensively investigated for a wide range of applications in cell biology ^13^. But, instead of merely measuring the cell stiffness at the surface (where Hertzian mechanics applies), or at a single fixed depth (where the Sneddon model is typically relevant), the modulus is determined as a function of depth for all 16,384 indentation curves per dataset. The position of the specimen surface must first be determined for each indentation curve, based on the distance traversed by the tip until a common repulsive force is reached during the indentation approach. Although any contact force could be considered, a relatively large value of 4 nN is implemented here, sufficient that cell deformation around the AFM probe will occur as previously predicted^30, 31^ and directly observed^12^, confirming applicability of the Sneddon model.

Accordingly, Figure 2A presents a 40 μm by 40 μm, 128 by 128 pixel topographic image, calculated from the contact point of all 16,384 force-distance (‘Fd’) curves in a single DDIP experiment of living A2780 human ovarian carcinoma cells. The image reveals part of an A2780 cell protruding 4-6 μm from the surface, positioned in the bottom center of the image, as well as part of another cell protruding as high as 8-10 μm at the upper right. The remainder of the field of view appears to be planar, compatible with the glass substrate on which the cells were cultured.

**Figure 2.**
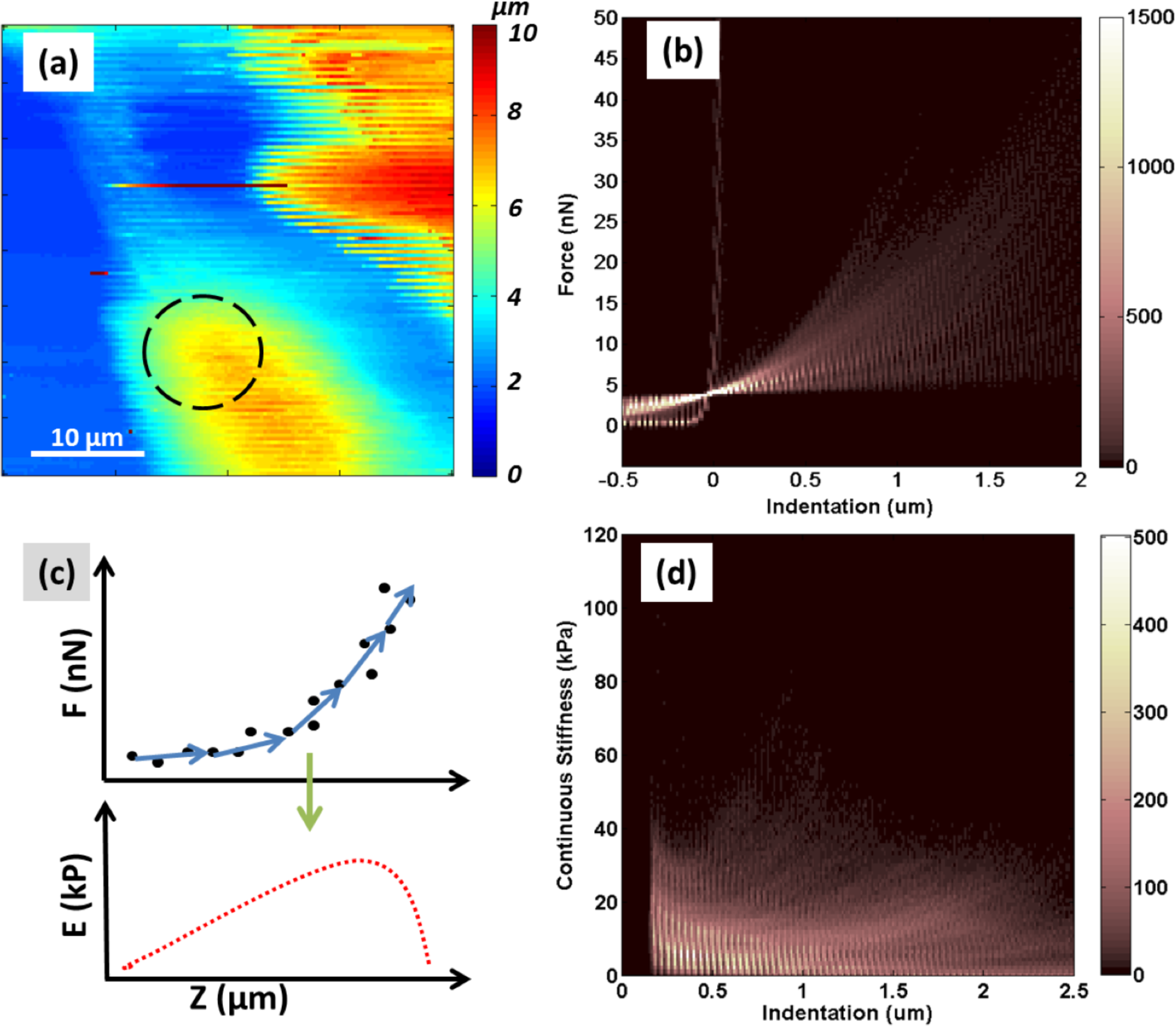
(a) Surface topography of A2780 cell created from 128x128 force curves. (b) Multidimensional histogram (counts indicated by color contrast) of all of the force (nN) vs. distance μm) (FD) curves. (c) As sketched, data from each force curve (location) is converted into a distinct continuous stiffness vs. indentation curve using Sneddon’s model. (d) Multidimensional histogram of continuous stiffness (kPa) vs. indentation μm) of the force volume data shown in Figure 2B.

To better explain the origin of the surface topography, it is useful to consider the corresponding indentation curves. While individual force spectra are often plotted in such circumstances, here a new approach is presented whereby a multi-dimensional histogram is generated of all of the 128*128 nanoindentations (color contrast indicates histogram counts), Figure 2B. This reveals three main indentation phenomena. 1) On contact (indentation depth = 0 μm), some force curves rise essentially instantaneously to the maximum force considered (50 nN). This can only result from tip interactions with a noncompliant surface relative to the spring constant of the AFM cantilever, in this case identifying the substrate as opposed to the substantially more compliant cells. 2) The force curve sometimes increases more slowly (5-40 nN/μm) down to an indentation depth of 0-2 μm, and then climbs abruptly as for the first case when the substrate was contacted. This suggests initial indentation into a compliant material such as a cell body or extra-cellular matrix, followed by interaction with the much stiffer underlying substrate. Note that such secondary abrupt rises in force are not apparent in the F-d histogram of Figure 2B, since there is not a single depth where this occurs throughout the imaged area. Considering individual force curves, or adjusting the contrast levels in the histogram, would reveal such events. 3) The repulsive force rises smoothly with indentation up to a depth of 2 μm (or beyond). Only data to 2 μm is considered here so that tip-sample interactions alone are considered; otherwise, cantilever-sample interactions can dominate once the indentation depth approaches the tip length (3 μm). Notably, for spectra from the compliant regions, a repulsive force is typically experienced up to 1-2 μm before the calculated 4 nN contact point, suggesting interactions between the tip and outlying cellular features that are regularly identified with optical microscopy but seldom seen for AFM of living cells (e.g. ruffles, cilia, or polysaccharide chains extending from the cell membrane). Such structures are dynamic on the time scale of these measurements (tens of minutes), though, essentially introducing noise into the images unless a relatively high contact threshold is employed as implemented here. This allows a more consistent focus on subsurface features as well, the primary objective of this work.

Based on the Sneddon equation, a least squares fit is performed for indentations beyond 80 nm for each force curve (Figure 2C, top). Notably, this is completed at a selectable number of depths (5 in the sketch), over fixed indentation ranges. This provides, for each measured location (pixel), a distinct continuous stiffness vs. indentation curve (Figure 2C, base). Based on several hundred distinct depth steps per force curve, Figure 2D presents a histogram of the resulting ‘continuous stiffness’ upon analyzing all 16,384 F-d curves (pixels). In most cases (>250 pixels at any given depth), the measured modulus is either <40 kPa (when contacting a cell or cell structures), or it is off-scale (i.e. interacting with the GPa-scale modulus of the glass coverslip). Throughout this work, as is common in AFM literature of compliant materials, stiffness and modulus are used interchangeably.

Analyzing these results further, indentation ranges of 0-80 and 80-500 nm are explicitly considered in Figure 3 (note log scales used for the contrast). For the first 80 nm, the Hertz model is used to map the contact stiffness (Figure 3A) and its corresponding standard deviation error is presented as a percentage of the calculated stiffness (Figure 3C). These elasticity maps for the A2780 cells up to 80 nm are then compared with the modulus (Figure 3B) and error (Figure 3D) calculated between 80 and 500 nm of indentation depth obeying the Sneddon model. For regions where the tip contacts the underlying glass coverslip during the indentation range considered, the calculated stiffness reaches GPa values. Such areas are denoted as NaN in the stiffness and error maps (black contrast) in order to focus on the mechanical properties of the cells instead of the invariant substrate.

**Figure 3.**
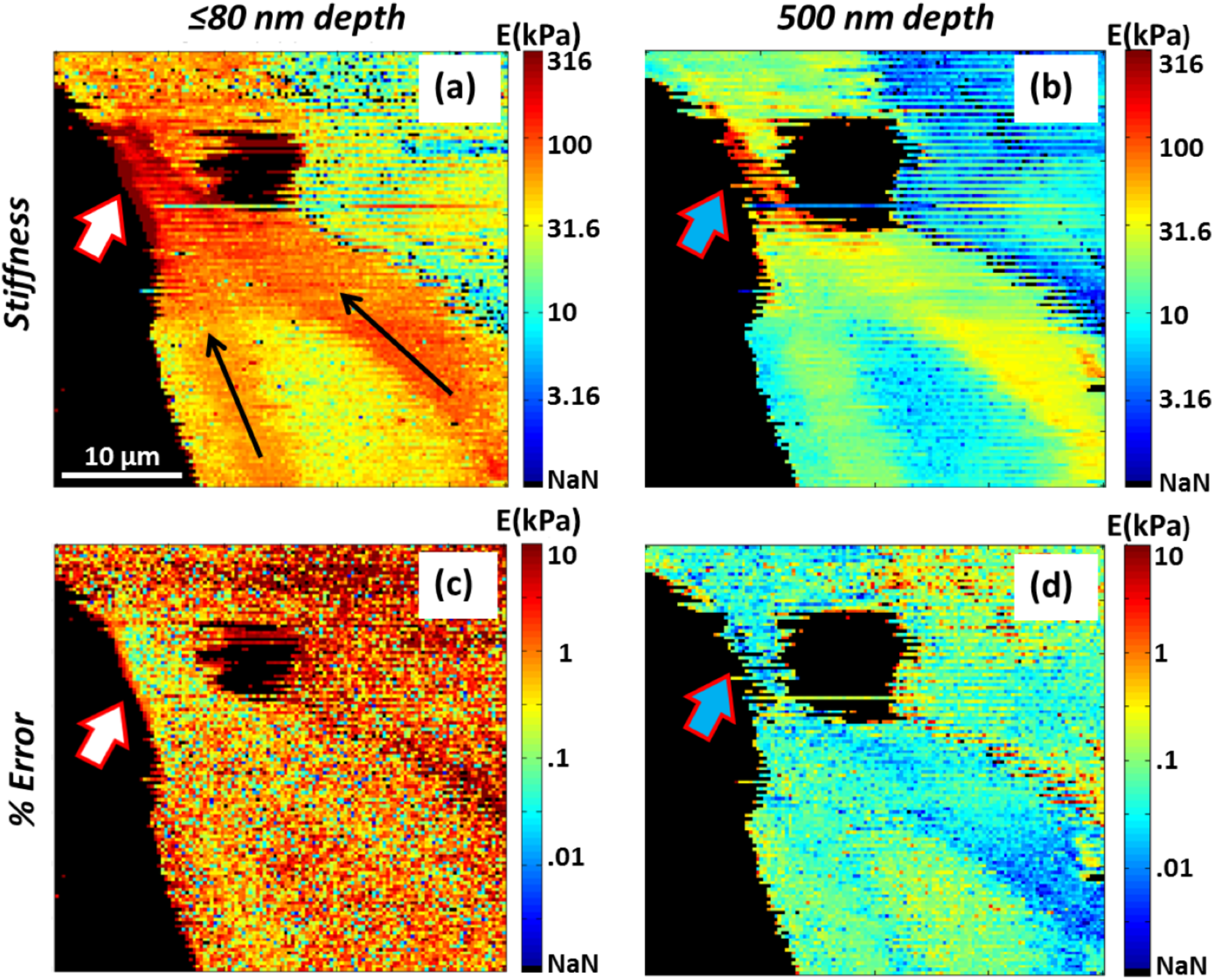
Calculated modulus and percentage standard deviation error maps of A2780 cells; note log scales to best present the range of measured features. (a) Indentation modulus up to 80 nm depth following Hertz mechanics. Overlain narrow black arrows indicate a probable extension of the cytoskeleton forming a pseudopodia marked by the short broad arrow. (b) Map of the modulus calculated according to Sneddon mechanics between 80 and 500 nm of indentation depth, with generally higher values due to incorporating a different mechanical model but overall similar trends in the mechanical properties. (c) Standard deviation of the calculated modulus, shown as a percentage error with respect to the locally calculated modulus up to 80 nm, is as high as 10% for most regions on the cell but drops to ~0.1% at the comparatively stiff pseudopoidal extension. (d) Corresponding percent error (standard deviation) of the Sneddon derived modulus for 500 nm indentations.

Long black arrows are sketched onto the 80 nm modulus map of Figure 3A to highlight regions of enhanced stiffness. When compared with the modulus map at 500 nm of indentation (Figure 3B), a similar pattern is observed though there is a difference in the modulus of almost an order of magnitude. It is hypothesized that these areas identify a high concentration of actin filaments that have developed along the sides and near the apical surface of the cell, bundling together to form a pseudopodial extension (block arrow) bridging toward the edge of another cell at the upper left of Figure 3. Such actin filaments tend to form extended networks within a cell body in response to stress, supporting structural protrusions to bear mechanical loads ^32^. DDIP therefore uniquely allows the stiffness of pseudopodia, actin filaments, and/or other cellular structures localized to certain depths to be 3-dimensionally investigated.

Further insight is available when the measured stiffness is correlated with maps of the error in the modulus calculations. As already described, these are determined based on the rms error of the regression fits from the raw Force-distance curves, Equations 6 and 7. For Figure 3C, this error is relatively uniformly distributed, with a speckle pattern indicating that it is dominated by position-independent noise in the data (note that the error is not calculated for pixels where the stiffness is off-scale, labelled as ‘NaN’). At the pseudopodia, however, the average stiffness error is approximately 1 order of magnitude smaller than in the rest of the cells. The same is true, but to a lesser degree, in the regions identified as being related to actin fibers, particularly apparent for the error map at the 500 nm probing depth (Figure 3D). One common explanation of such a locally enhanced stiffness is a possible convolution with the mechanical properties of the underlying substrate. However, in these locations the specimen is still 2-4 um thick according to Figure 2a. More significantly, substrate-induced artifacts should make the fit to the mechanical model (and hence the error) worse, whereas the error maps conversely reveal the opposite. This suggests that the local modulus is simply better modeled by elastic mechanics (particularly Sneddon) in these stiffer, actin-filament-rich regions.

Although a primary objective of DDIP is to identify depth-dependent mechanical properties, simply comparing the magnitudes of the modulus and modulus error at the distinct 80 and 500 nm depths depicted in Figure 3 does not provide any obvious evidence of distinct sub-surface features. The magnitude is different, but the relative stiffness is similar throughout the field of view. This is also true for stiffness maps based on the entire force curve at each pixel, which is the most common analysis of AFM-indentation measurements (not shown for brevity). However, since DDIP calculates the stiffness continuously over all depths, a promising approach is to determine the peak stiffness (Figure 4a). This is particularly powerful when considered alongside the depth at which this peak stiffness occurs (Figure 4b). As an example, the dashed circular regions sketched on the figures identify a roughly 50 μm^2^ area. This area exhibits a relatively higher stiffness than the surroundings, with an average of ~70 kPa compared to ~30 kPa nearby. This occurs at a depth of approximately 1.4 μm (Figure 4B) as compared to 1.8 μm. The cell thickness in this area is still 5 to 6 μm, though, so again substrate-related artifacts are ruled out. Instead, this zone of high stiffness can only be explained by sub-surface cellular sub-structures. It may be caused by unknown features suspended in the cytoplasm of this human ovarian carcinoma, but based on the location it is likely either a cross-over point of cytoskeletal fiber bundles, and/or a region with relatively stronger ties to focal adhesion points, pseudopodial extensions, etc. This uncertainty could be addressed in future work by combining DDIP with simultaneous optical fluorescence studies.

**Figure 4.**
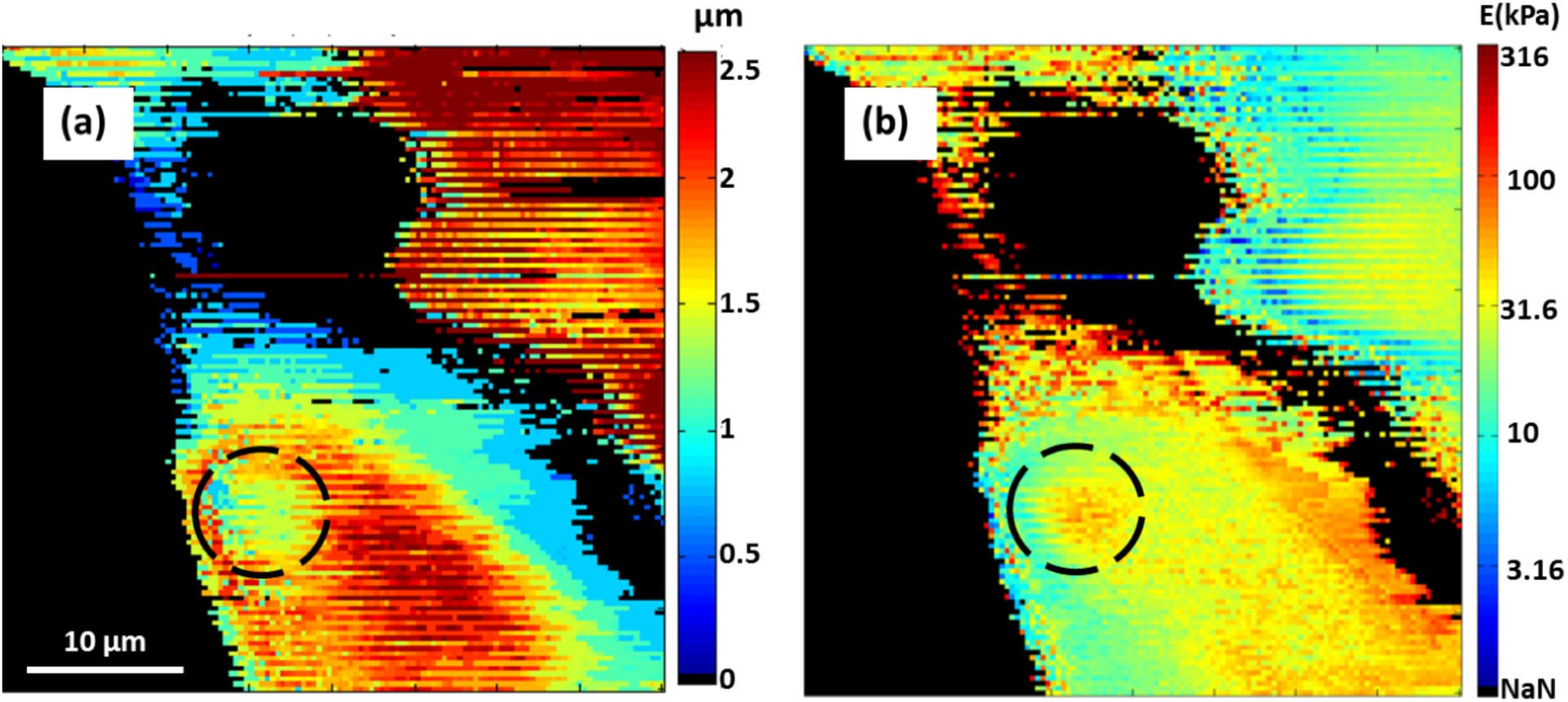
(a) Map of the peak modulus experienced at any depth for each pixel from Figures 2 and 3, excepting substrate effects. Note a log scale is used to best present the range of modulii, with ‘NaN’ identifying the many orders of magnitude stiffer substrate. (b) The height at which the maximum modulus occurs.

### Modulus and error mapping of Hydrogel

In order to further test this analytical approach, indentation measurements were made with 0.5% and 1% agarose gel as a homogeneous proxy for general cellular mechanical measurements. For each specimen, an array of 16x16 indentations was performed over a 20 μm by 20 μm area, repeated 3 times in different locations on the gels. Images similar to Figures 2-4 result, but are spatially featureless. Therefore, the results for this experiment are more meaningfully presented as histograms of the calculated modulus (Figure 6) determined at each of three distinct depths: 50 nm according to the Hertzian model (1^st^ row), 500 nm according to the Sneddon model (2^nd^ row), and 1 micron also obeying Sneddon mechanics (last row). The percentage error for each calculation is shown as well (right column), based on the standard deviation of the fit to each model as with Figures 3c-d. Note that each point on the plots represents the mean value from the three repeated experiments for each Agarose gel, with error bars indicated the standard deviation.

**Figure 5.**
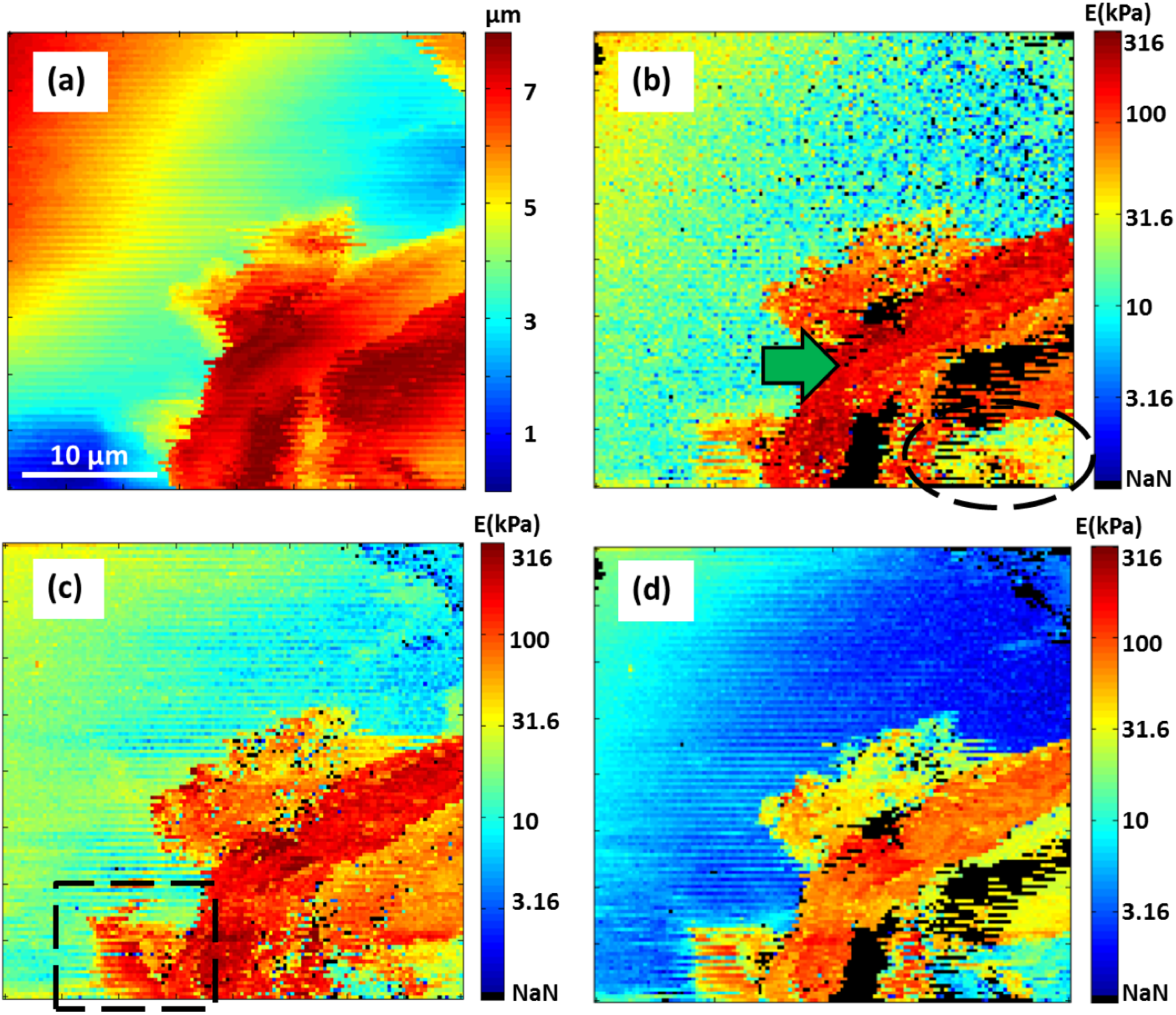
(a) Topography of NIH3T3 cell and (b) DDIP generated modulus map assuming Hertz model for indentation less than 80 nm. The broad arrow marks a proposed Circular Dorsal Ruffle (CDR); the black dashed oval at bottom right highlights a similar modulus inside the F-actin ring as in the rest of the cell body as seen in the upper half of the image. (c) DDIP derived modulus map based on Sneddon mechanics at 250 nm depth. The square dashed rectangle at bottom left reveals a feature that is not visible in topography and is only partially apparent at 80 nm. (d) There is an overall decrease in stiffness for the Sneddon derived modulus 500 nm beneath the surface as compared to 250 nm or shallower, though there are several regions with the opposite behavior as highlighted by the higher stiffness band extending from the overlain broad arrow.

**Figure 6.**
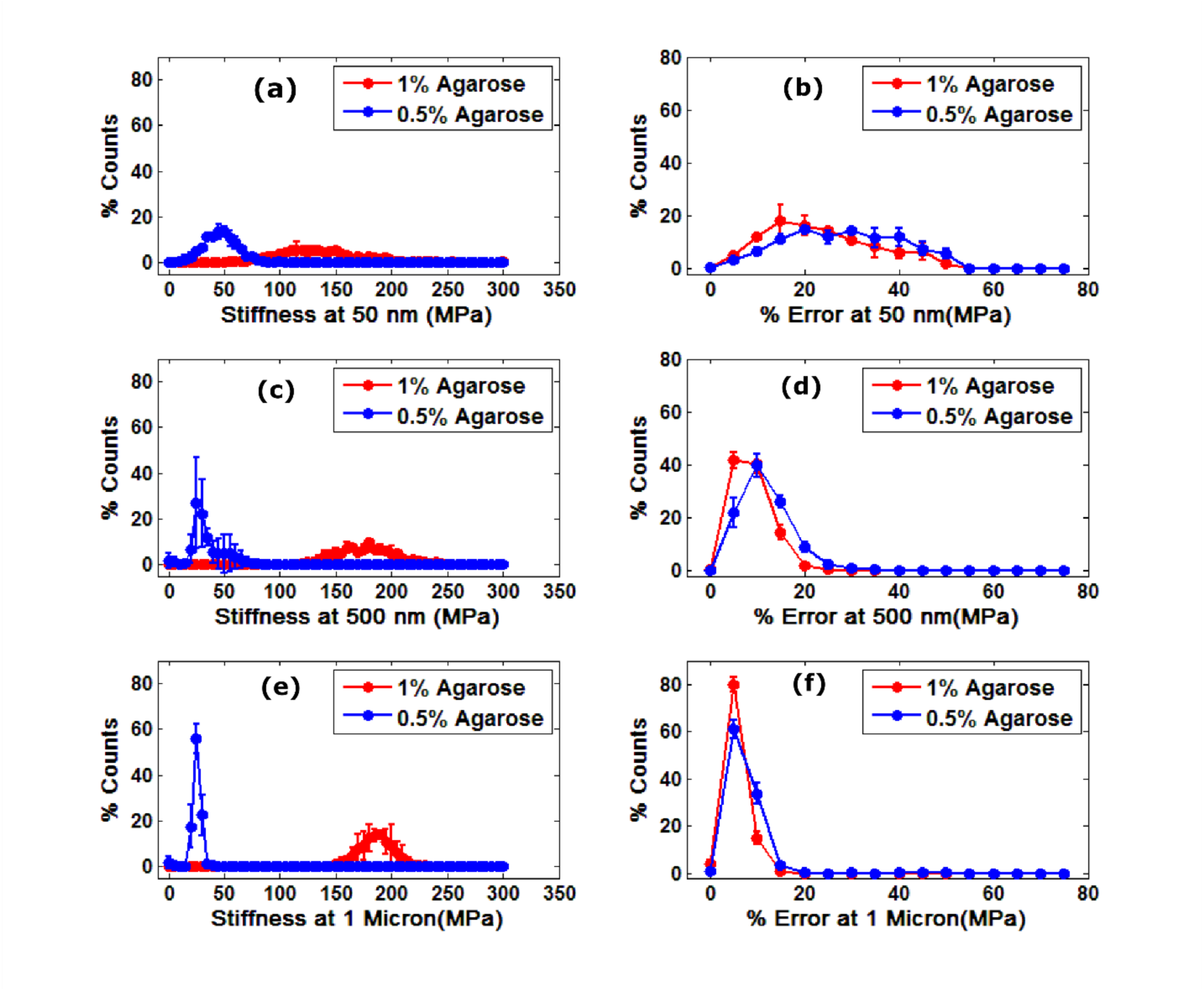
Modulus and percentage error histograms of 0.5% and 1% agarose gels at several depths (each point presents the average and standard deviation from 3 gels, with 16×16 indentations per gel). (a) Hertzian derived moduli at 50 nm depth and (b) their error reveal a wide distribution of measured moduli, and error, for both 0.5 and 1% agarose samples. (c) Sneddon derived moduli at 500 nm and (d) their error exhibit substantially narrower distributions, with less average error, as compared to shallow indentation (50 nm) data. (e) Sneddon derived moduli at 1 micron depths and (f) corresponding errors are sharper still. Note the gradual apparent decrease in modulus for 0.5% Agarose, and conversely a gradual increase for 1.0% agarose, with depth, independent of error in the modulus calculations.

The first observation for the gels is that with increasing depth (top to bottom row), the distribution of calculated moduli tightens as the depth increases from 50 nm to 1 μm, independent of the gel concentration. The corresponding error calculations exhibit the same trend, which was also generally exhibited by the A2780 cells in Figures 3c-d. For the deepest stiffness calculations the mean error in the fit is roughly 5% with a maximum of roughly 15%, whereas the mean is closer to 15% ranging as high as 25% just 500 nm beneath the surface, and even worse for Hertzian calculations at 50 nm depth. Based on such percentage error measurements, seldom reported in the AFM literature, this leads to two possible conclusions. The gel may be inhomogeneous for indentations on the order of 50 nm, causing a wide range of measured local moduli for multiple measurements which is averaged out only for deeper and correspondingly broader indentations that simultaneously interact with the entire range of heterogeneities. This is unlikely, however, as it would suggest a pore structure and/or polymerization inhomogeneities at ~50 nm scale are much larger than anticipated for agarose gels^33^. More reasonably, at least for homogeneous structures with bulk moduli in the 10 to several hundred MPa range given the tip/sample geometry in the AFM and the loading conditions, Sneddon calculations at depths on the order of 1 um are simply more representative of the actual mechanical contact than Hertzian models at depths in the tens of nanometers.

The second observation from Figure 6 is that the calculated modulus changes with depth. The peak in the apparent stiffness of the 1% Agarose gel (left column) increases slightly with depth, while the stiffness for the initially more compliant 0.5% gel conversely reduces by up to a factor of 2. The divergence of the stiffness with depth depending on the Agarose concentration supports that these results indicate real changes in properties instead of possible artifacts introduced by the experiment or analysis (which would not diverge). It also cannot be attributed to the change of model from Hertzian to Sneddon since the trend continues within the Sneddon approximation for depths of 500 nm to 1 μm. Instead, the results reveal an enhanced stiffness or ‘crust’ at the surface of the 0.5% agarose, and the opposite for the 1% agarose. This is most likely caused by changes in the porosity, crosslinking, and/or water content of the hydrogels with depth, however unexpected given that the specimens were prepared identically aside from the agarose concentration. Being able to compare calculated moduli, and error, as a function of depth strengthens such conclusions, confirming the value of DDIP and proper error analysis for identifying mechanical heterogeneities in 3-dimensions for compliant materials.

On 0.5% agarose this reduction in stiffness from 50 nm to 1 micron is by factor of 2 whereas on live cells it’s greater than 3. CDRs are bundled actin filaments and as discussed in introduction, cytoskeleton can effectively transfer load from local point of stress to focal adhesion centers at the base of the cells. Hence, there is an instantaneous decrease in stiffness which is larger as compared to data collected on 0.5% agarose. This dissipation of load on cell highlights the efficacy of whole cytoplasmic network. Rest of the cell body shows response similar to what we have observed in 0.5% agarose gel. Based on the tensegrity model that defines intracellular mechanical properties of cytoskeletal network, DDIP gives us a possibility to test different kinds cytoskeletal fibers, actin filaments, intermediate filaments and microtubules^34^. These fibers that are always present in pre-stressed state and with depth wise study one can focus one particular kind of cytoskeletal element (CE) and measure how it is responding to mechanical stimulus from an AFM probe^21^. Cellular tensegrity and mechnotransduction go hand in hand for cells to respond immediately to any external stress. This intricate network of cytoskeletal filaments and mechanoreceptors that helps a cell to develop its prosthesis according the changes in its environment^21, 35^. To segregate CE with respect to their role in external load dissipation, AFM has to be coupled with other detection methodologies like fluorescence to simultaneously record any visual changes during indentation measurements. In one such study mitochondria was fluorescently labelled and its displacement with respect to mechanical forces was measured. Mitochondria are transported within the cell via tubular network^13^. Application of mechanical force results in displacement of mitochondrion or similar organelles within the cell. Degree of displacement is affected by presence tubular network which in turn affects porosity in its immediate surroundings.

### NIH3T3 Mouse Fibroblasts

Finally, DDIP was employed with live NIH3T3 cells, revealing 3-dimensional nanomechanics of a dynamic cellular structure known as a Circular Dorsal Ruffle (CDR) which has not previously been reported in AFM literature. Figure 5a displays a topography map akin to that in Figure 2a, only in this case the underlying substrate is not exposed. Instead, part of a compliant cell with a smoothly sloping topography fills the field of view, with an up to 5 μm protrusion at lower right covering approximately one-fourth of image. For comparison, simultaneously acquired DDIP-generated modulus maps are shown for indentation depths of 80 nm according to Hertzian mechanics (Figure 5B), 250 nm following the Sneddon model (C), and 500 nm according to Sneddon (D). At the edge of the topographic protrusion, a ring of enhanced stiffness ranging from 30-300 kPa is apparent depending on the depth, marked by the broad arrow in (b). For greater depths, this ring is still detected but is more compliant, suggesting that the feature is present at or near the apical surface of the cell.

It is proposed that this ring-shaped feature is a circular dorsal ruffle (CDR). CDR’s are F-Actin rich membrane projections with extensive bundling and branching of actin fibers. Therefore they can be expected to exhibit a higher stiffness than actin fibrils or other more compliant structures within a cell body ^36^, compatible with the ~300 kPa modulus determined here. It is only during formation of circular dorsal ruffles (CDR), however, that actin filaments assemble into ring-shaped structures as observed. The rate of turnover for CDR’s varies with cell type, but they have been reported in NIH3T3 cells to last up to 60 mins according to optical imaging^32, 36, 37^. In live cell nanomechanical probing, however, CDR’s have not previously been reported.

Although entire CDR’s were not imaged, the dashed oval in the bottom right corner of Figure 5b reveals a modulus inside the ring-shaped structure similar to that outside the ring, as expected for CDR’s which primarily protrude normal to the surface of the cell by as much as 10 μm ^38^ as also observed here. Uniquely available through DDIP, occasional bands of locally enhanced stiffness (laterally and in depth) are also observed within the CDR. This is exemplified by the approximately 3 μm by 8 μm region of stronger contrast 500 nm beneath the surface, identified by the block arrow in Figure 5d. A similar sub-surface mechanical enhancement, not apparent in topography and barely suggested in indentations at 80 nm or 500 nm, is resolved at a depth of 250 nm within the overlain dashed rectangle (Figure 5c).

Considering sub-surface nanomechanics one step further, the continuous stiffness as in Figure 2d is shown in Figure 7 for a single row from the 128x128 pixel DDIP dataset used for Figure 5. Specifically, the Sneddon-derived modulus is determined centered at depths varying from 200 nm to 1 μm. 2 populations of pixels are compared. Solid lines present the continuous stiffness for 40 pixels along a single scan row within the standard cell body, located to the left of the CDR as identified by the overlain line in the inset image (which is otherwise identical to Figure 5b). Dotted/dashed lines identify the continuous stiffness for 40 pixels within the CDR, located in the right ~1/3^rd^ of the image along the same scan row also as indicated in the inset. The stiffness profiles are dramatically different for these two specimen regions, as expected for the compliant cell body (~5-10 kPa) compared to localized actin structures (~20-100 kPa) characteristic of CDR’s.

**Figure 7.**
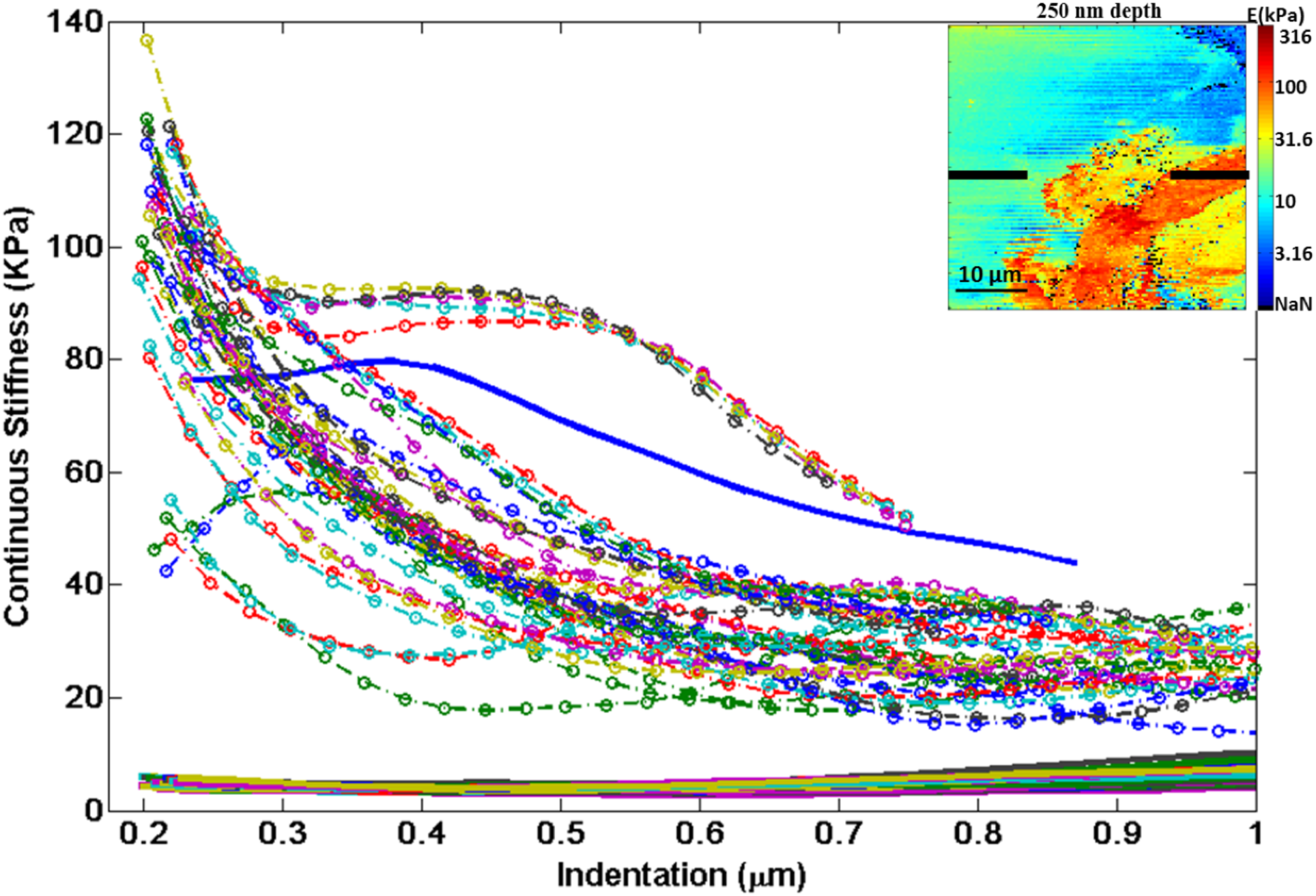
Continuous Stiffness vs Indentation curves up to a depth of 1 micron. The curves are extracted from along the dotted line marked in the modulus map from a 250 nm depth (Figure 5c). The first 40 curves (marked -) show no gradual increase or decrease in stiffness because the probe is indenting into the cell body with no appreciable actin in the vicinity. In the last 40 curves (marked–o.), the probe is indenting into the proposed CDR structure, where the continuous stiffness generally decreases but also frequently increases as a result of sub-surface structure.

Within the CDR, two distinct depth-dependencies can be distinguished. Type 1 behavior, at most locations within the CDR, exhibits an initially strong calculated modulus, which diminishes smoothly with indentation. For type 2, a few regions exhibit peaks or at least plateaus in the modulus over depth ranges of several hundred nm. As with a few regions identified in the discussion of Figure 5, this suggests localized, sub-surface mechanical structures.

Overall, it is important to note that for all cell measurements, the calculated moduli near the surface can be relatively stronger than several hundreds of nm beneath the surface or deeper (as with Figure 2d, and type 2 regions in Figure 7). Again, where this occurs in this work it is interpreted as a near-surface-enhancement of the local stiffness. Another possibility that must be considered, however, is essentially hydrodynamic: as the AFM probe used in these experiments begins to indent the cell, the pressure can rise rapidly with indentation, further complicated by necessary reconfiguration of the cell membrane and nearby features to accommodate the perturbing AFM tip. Force curves were collected at a scan rate of up to 32 μm/s, such that the first 200 nm of indentation occurs in as little as ~6 ms. This is faster than the time scale for water redistribution in some cells according to studies of the influence of cytoplasmic fluids on force-relaxation at short times scales near the apical surfaces of cells ^15^. Those studies used colloidal probes which applied an even lower pressure than the conical probes employed here ^39^. Future DDIP studies as a function of loading rate are therefore worthwhile to identify whether such nanomechanical enhancements near the cell surface are hydrodynamic or related to cytoskeletal structures, intracellular tubular networks, etc. If the latter, DDIP can enable novel investigations of variations in such surface effects for specific cell types, disease states, etc. But the strongly varied behavior of the modulus as a function of depth within a single cell, as well as the divergence of the modulus with depth for agarose hydrogel, confirms that the general observations provided by DDIP herein are not dominated by hydrodynamic or calculation induced artifacts, and instead uniquely reveal real specimen features that vary with position and depth.

## Conclusion

Systematic measurements of the depth dependence of the local modulus, and error, can help us to understand dissipation of local stress as a function of depth. This gives us a cut-off depth and error to accurately describe mechanical property of a single isolated cell or a tissue. We can also differentiate between stiffness of various pseudopodial extensions by calculating stiffness at different depths. This novel approach towards stiffness profiling can used to study membranal protrusions from cells apical surface like Villi or CDRs. In tissue, one can comment on mechanical properties of interconnecting boundaries between different cell and how extracellular matrix influences such properties. With ever expanding world of polymer research especially biopolymer research, here we present a strategy to check uniformity in polymerization up to few nanometer to a micron into the surface. We are moving into an age of synthetic cellulo-mechano platforms. By understanding different rheological parameters of cells under different synthetic environments can help us to develop more controlled artificial living systems. Above described analytical approach can provide answers to number of such nanomechanical properties.

## Author Contributions

All authors have given approval to the final version of the manuscript.

## Notes

Authors declare no competing financial interests.

## Graphical Abstract

**Dynamic and Depth Dependent Nanomechanical Properties of Dorsal Ruffles in Live Cells and Biopolymeric Hydrogels**

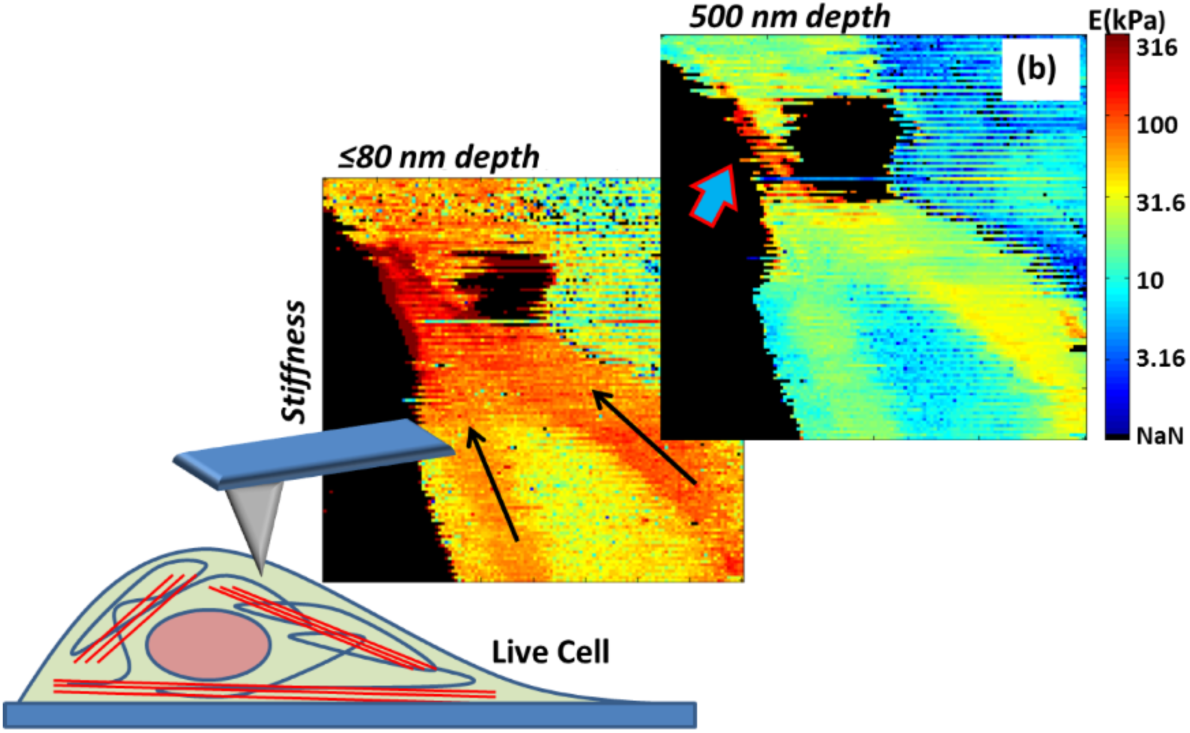

Varun Vyas^1 (✉)^, Melani Solomon^23^, Gerard G.M. D'Souza^2^, Bryan D. Huey^1 (✉)^

**Calculating the depth dependent modulus in live cells with Atomic Force Microscopy:** Atomic Force Microscopy investigations of live cells and biopolymers reveal fluctuations in stiffness as function of depth. Such tools have applications towards understanding sub-surface nanomechanical properties in various tissues and cell lines as well as how porosity of biopolymers is affected by differential hardening or polymerization of different components in a polymer. Here, depth dependent modulus of A2780 & NIH3T3 cell lines and Agarose was measured using pyramidal AFM probes.

